# Genomic diversity and antimicrobial resistance among non-typhoidal *Salmonella* associated with human disease in The Gambia

**DOI:** 10.1101/2020.10.30.316588

**Authors:** Saffiatou Darboe, Richard S. Bradbury, Jody Phelan, Abdoulie Kanteh, Abdul-Khalie Muhammad, Archibald Worwui, Shangxin Yang, Davis Nwakanma, Blanca Perez-Sepulveda, Samuel Kariuki, Brenda Kwambana-Adams, Martin Antonio

## Abstract

Non-typhoidal *Salmonella* associated with multidrug resistance cause invasive disease in sub-Saharan African. Specific lineages of serovars *S*. Typhimurium and *S*. Enteritidis are implicated. We characterised the genomic diversity of 100 clinical Non-typhoidal *Salmonella* collected from 93 patients in 2001 from the eastern and 2006 to 2018 in the western regions of The Gambia respectively. Phenotypic susceptibility applied Kirby Baur disk diffusion and whole genome sequencing utilized Illumina platforms. The predominant serovars were *S.* Typhimurium ST19 (31/100) and *S.* Enteritidis ST11 (18/100) restricted to invasive disease with the notable absence of *S.* Typhimurium ST313. Phylogenetic analysis performed in the context of 495 African strains from the European Nucleotide Archive confirmed the presence of the *S*. Enteritidis virulent epidemic invasive multidrug resistant West African clade. Multidrug resistance including chloramphenicol and azithromycin has emerged among the West African *S.* Enteritidis clade 7/9 (78%) with potential for spread, thus having important implications for patient management warranting systematic surveillance and epidemiologic investigations to inform control.

**Data summary:** Sequences are deposited in the NCBI sequence reads archive (SRA) under BioProject ID:PRJEB38968. The genomic assemblies are available for download from the European Nucleotide Archive (ENA): http://www.ebi.ac.uk/ena/data/view/. Accession numbers SAMEA6991082 to SAME6991180

## Introduction

Non-typhoidal *Salmonella* (NTS) serovars are a common cause of foodborne gastroenteritis but can also cause severe disseminated infections dependent on the pathogen’s virulence and the host’s immune status [1,2]. Globally, there are over 2,800 serovars, some of which are adapted to non-human hosts [3]. The global annual estimate of *Salmonella* gastroenteritis is 93.8 million illnesses with 155,000 deaths [4]. The highest mortality occurs in Africa which accounts for 4,100 deaths annually with an incidence of 320/100,000 population [4]. Most cases of *Salmonella* gastroenteritis in immunocompetent hosts are self-limiting and do not require antimicrobial therapy; however, infections in infants, the elderly and immunocompromised patients do require antimicrobial treatments such as ciprofloxacin, 3^rd^ generation cephalosporins or azithromycin [5].

The clinical characteristics of non-typhoidal salmonellosis emerging in Africa represent a changing disease pattern, from gastroenteritis to invasive disease with a case fatality ratio of 20-25% [1,6–8]. Although NTS in diarrhoea is less well characterised in Africa, it may be predisposition to invasive disease. Invasive NTS (iNTS) disease, mainly bacteraemia. iNTS is globally estimated at 3.4 million illnesses annually, disproportionately affecting those in sub-Saharan Africa (sSA), with over 50% associated with HIV infection, malnutrition, recent malaria and children between 6 months to 3 years of age [6,8,9]. iNTS disease has a markedly different presentation, closer to enteric fever in its clinical form than typical NTS disease [8]. The predominant serovars responsible for invasive disease in sSA are specialised lineages of *Salmonella enterica* serovars Enteritidis and Typhimurium that are distinct from those circulating in other parts of the world [1,10]. Whole genome sequencing (WGS) has provided new insights into the host adapted signatures associated with pathogenicity and metabolism of these *S.* Typhimurium lineages and *S.* Enteritidis clades characterised by genomic degradation and accessory genome [1,11].

The closely related *S.* Typhimurium ST313 lineages I and II evolved independently around 52 and 35 years ago respectively with the acquisition of the chloramphenicol acetyltranferase (*cat)* resistance gene [7]. Both lineages have been shown to carry a *S.* Typhimurium virulence plasmid, commonly known as pSLT, which also encodes genes conferring resistance to common antimicrobials including tetracycline, sulfamethoxazole-trimethoprim and chloramphenicol [7,11]. In addition, two related, but phylogenetically different epidemic clades of *S.* Enteritidis ST11, the West African clade and the Central/Eastern African clade, characterised by the presence of chloramphenicol acetyl resistance genes *catA1* and *catA2*, respectively, plus the incomplete set of *tra* genes, emerged between 1933 and 1945 [1]. The utility of second-line antimicrobials such as fluoroquinolones, azithromycin and extended spectrum cephalosporins is limited in treating these emerging MDR strains [12].

In The Gambia, iNTS remains a leading cause of invasive diseases, [13] unlike *S.* Typhi which causes typhoid fever in many low and middle income countries (LMIC). We previously described regional serovar variation and emerging MDR in The Gambia [14,15]: *S.* Typhimurium was found to be more prevalent in the western region, while *S.* Enteritidis, including MDR strains, were found to be more prevalent in the eastern region. In this context, we performed whole-genome analysis of clinical NTS isolates to determine prevalent genotypes and antimicrobial resistance genes. The resulting analysis can be used to help guide clinical management and control of NTS diseases in The Gambia.

## Methods and Materials

### Study setting and population

The study was conducted at the Medical Research Council Unit The Gambia at the London School of Hygiene and Tropical Medicine (MRCG @ LSHTM) using clinical NTS isolates from the eastern (Upper River Region) and western (West Coast Region and Greater Banjul Area) regions of The Gambia (Figure 1). The eastern region, located on the far east side of the river Gambia, is the commercial centre and a busy economic hub, with an estimated population of 200,000 people. It is an important transit point for merchandise and people going into eastern Senegal, Mali and Guinea Conakry. The western region is densely populated, with a population of over 1 million people including the capital city, Banjul (Figure 1) [16]. Malaria declined in recent years but remains endemic with peak transmission occurring from July to November [17]. Malnutrition remains a problem with the prevalence of underweight, stunting and wasting among children under 5 years old estimated at 16.4%, 25.0% and 4.3% respectively [18]; HIV prevalence among adults aged 15 – 49 years is estimated at 2.1% [19].

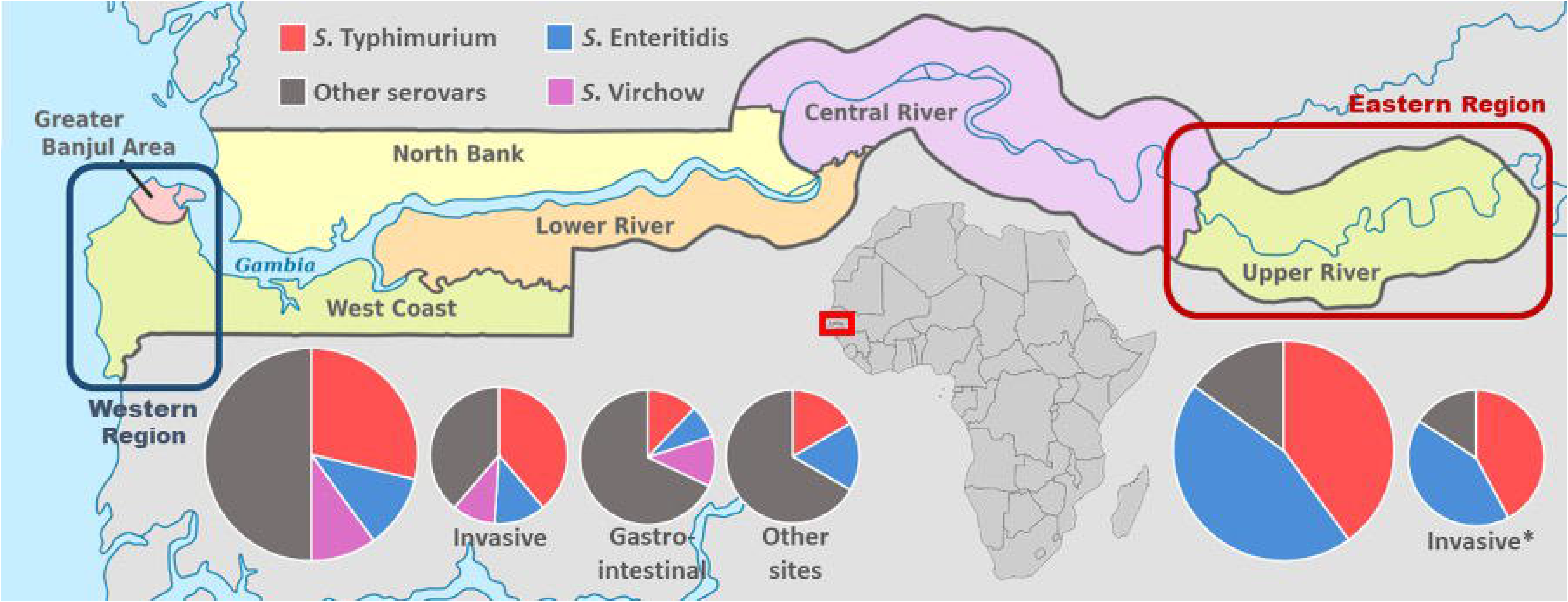

### Sample collection, microbiological procedures and antimicrobial susceptibility testing

The study evaluated 100 clinical NTS from 93 patients admitted to hospital with suspected sepsis, gastroenteritis or other focal infections in the eastern and western regions of The Gambia (Table 1). Seven patients had multiple samples collected during the same infection episode (Table 2). Three patients had concurrent bacteraemia and gastroenteritis, two had bacteraemia with meningitis whilst two had bacteraemia with two sampling episodes. All NTS from the eastern region (20) were isolated in 2001 from 18 patients, and those from the western region (80) were isolated between 2006 to 2018 from 75 patients (Table 1). All isolates were stored in 15% (v/v) glycerol broth at −70°C. The isolates were grown on MacConkey agar overnight at 37°C in the Clinical Microbiology Laboratory. The laboratory is accredited to Good Clinical Laboratory Practice (GCLP; 2010) and ISO15189 (2015) as previously described [14]. Antimicrobial susceptibility for amoxicillin-clavulanate, ampicillin, cefotaxime, cefoxitin, ceftazidime, ceftriaxone, cefuroxime, chloramphenicol, ciprofloxacin, gentamicin, nalidixic acid, sulfamethoxazole-trimethoprim and tetracycline, were tested on Mueller-Hinton agar (MHA) using the Kirby-Bauer disk diffusion method. Interpretation was done according to the 2017 Clinical Laboratory Standard Institute (CLSI) guidelines [20]. Antimicrobial agents were from BD Oxoid (Basingstoke, United Kingdom) and *Escherichia coli* (ATCC 25922) was used for quality control.

**Table 1.** Baseline characteristics of Gambian non-typhoidal *Salmonella* disease patients from whom isolates were cultured for use in this study.

|  |  | *N (%)* | *eastern region* | *western region* |
| --- | --- | --- | --- | --- |
| *Age range* | 0-4 years | 42 (45.2) | 4 (25.0) | 38 (50.7) |
|  | 5-14 years | 21 (22.6) | 12 (65.0) | 9 (12.0) |
|  | ≥15 years | 26 (27.9) | 2 (10.0) | 24 (32.0) |
|  | Unknown | 4 (4.3) | 0 | 4 (5.3) |
| *Gender* | Male | 51(54.3) | 10 (55.6) | 41 (54.7) |
|  | Female | 38(41.5) | 7 (38.9) | 31 (41.3) |
|  | Unknown | 4 (4.2) | 1 (5.5) | 3 (4.0) |
| *Source* | Invasive disease | 68 (68.0) | 19 (95.0) | 49 (61.2) |
|  | Gastroenteritis | 26 (26.0) | 1 (5.0) | 25 (31.3) |
|  | Other | 6 (6) | 0 | 6 (7.5) |
| *Serovars* | *S. Enteritidis* | 18 (18.0) | 9 (45.0) | 9 (11.2) |
|  | *S. Typhimurium* | 31 (31.0) | 8 (40.0) | 23 (28.8) |
|  | *S. Virchow* | 8 (8.0) | 0 | 8 (10.0) |
|  | Other serovars* | 43 (43.0) | 3 (15.0) | 40 (50.0) |

\*Shown in figure supplementary table

**Table 2.**
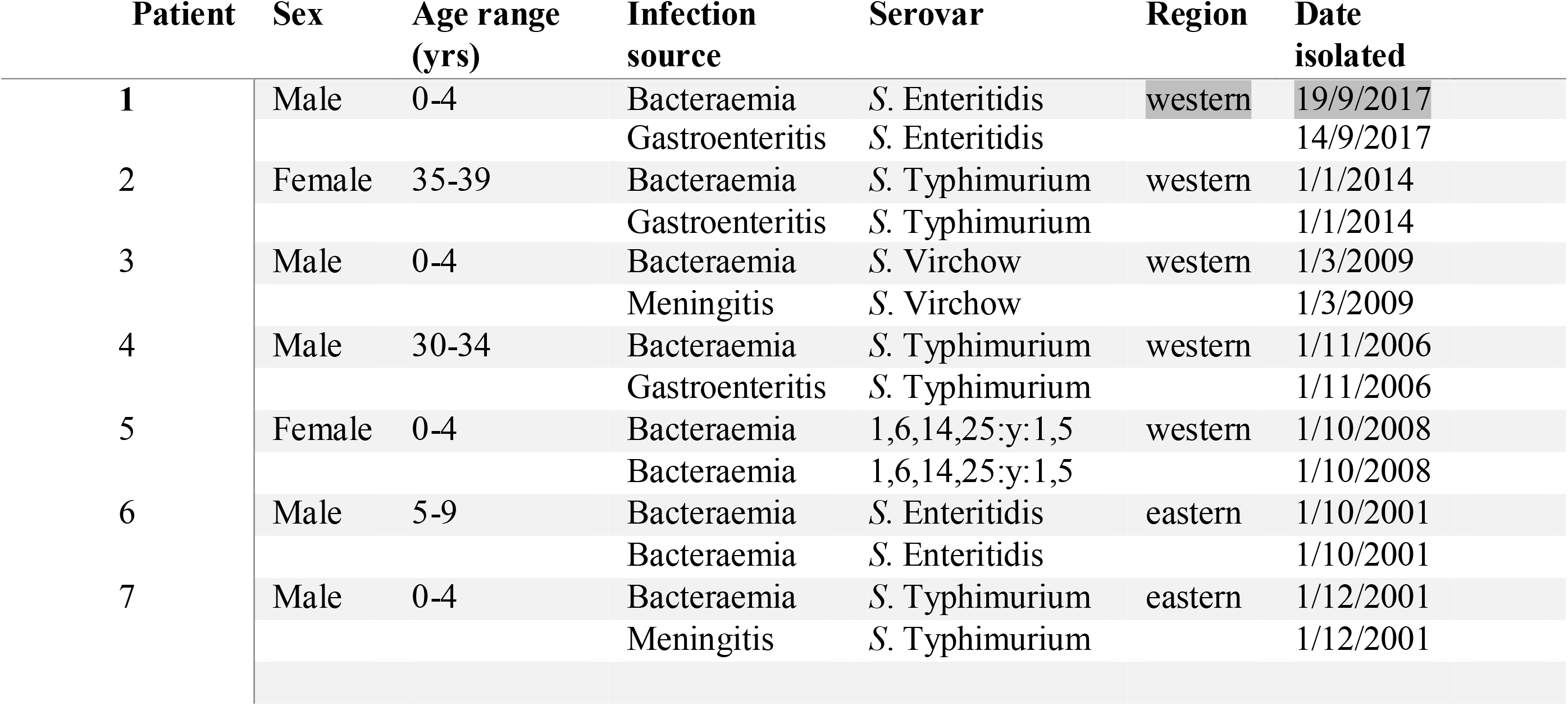
Patients with multisite NTS simultaneous infections.

### DNA extraction and whole genome sequencing

Genomic DNA was extracted and sequenced in two locations; the MRCG (n=33), and sample processing (n=67) at the University of Liverpool (UK) with DNA sequencing at the Earlham Institute (UK). The extraction protocol for the MRCG used the QIAamp DNA Mini kit (Qiagen, Germany) to extract the DNA from 1mL of an overnight culture grown in triple soya broth (TSB) from BD Oxoid (Basingstoke, United Kingdom) at 37°C, according to manufacturer’s instructions, and quantified using a Qubit fluorometer (ThermoFisher, Qubit dsDNA HS Assay). Libraries were prepared using the Nextera XT kit using the Illumina MiSeq system. The DNA extraction and sequencing of samples processed at the University of Liverpool were carried out using an optimised method for large-scale sequencing [21], including the bespoke LITE (Low Input, Transposase Enabled) pipeline for library construction, and Illumina HiSeq sequencing technology. Both sites used the 2×150 bp read protocol.

### Genome assembly and *in silico* analysis

The quality of the raw reads was assessed using FASTQC [22] where on average, all reads had a quality Phred score (Qscore) above 30. Paired-end reads were trimmed using Trimommatic [23] and assembled into contigs using SPades, with default settings [24]. *In silico* serotyping of the core genome MLST (cgMLST) and serovar was predicted using the *Salmonella in Silico* Typing resource (SISTR) platform [25]. eBurst Groups (eBGs) were assigned using the Enterobase platform which is based on the allelic identity that accounts for homologous recombination, defined by Achtman as closely related natural genetic clusters/populations of two or more sequence types connected by pair-wise identity or single locus variants [26].

### Phylogenetic analysis

Assembled contigs were annotated using Prokka (v1.14.6). The pan-genome was determined using Roary (v3.13.0) [27], taking the GFF files from Prokka as input with default settings. The pan-genome was aligned using Mafft (v7.464) to generate a high-quality sequence alignment. The alignment was used to create a maximum likelihood phylogeny using IQ-TREE (v1.6.12). The phylogenetic tree was visualised and annotated using the interactive Tree Of Life (iTOL). ITOL annotations input files for the tree were generated using custom python scripts (https://github.com/jodyphelan/itol-config-generators). Antimicrobial resistance (AMR), plasmids and virulence genes were detected by ResFinder [28], PlasmidFinder [29] and virulence factor gene database (VFDB) with a minimum coverage and nucleotide identity of 90% as the cut-off. Publicly available data was downloaded from the European Nucleotide Archive (ENA) to compare the strains from this study against other African strains. All sequence reads from the African samples belonging to the taxid 149539 (NCBI:txid149539) were downloaded and assembled/annotated using the same methods as detailed above. The pangenome phylogeny was constructed using the same methods as outlined above.

### Statistical analysis

The independent variables were serovars and the dependent variable was disease category. The relationships between the serovar and disease were analysed using logistic regression with measures of association expressed in odds ratios. No power calculations were performed and an alpha value of 0.1 was considered statistically significant. All statistical analyses were performed in Stata, Version 13.1 (StataCorp. 2013. Stata Statistical Software: Release 13. College Station, TX: StataCorp LP.)

### Ethical Review and Approval

The study received ethical approval from the Joint MRC/Gambia Government Ethics Committee (SCC1498).

## Results

### Isolate source, associated disease syndrome and regional serovars differences

One hundred isolates recovered from clinical samples from eastern (n=20) and western (n=80) regions of The Gambia were analyzed (Table 1). Isolates were recovered from patients of all ages, with a median age range of 5-14 years. Isolates from the eastern region were predominantly from invasive disease (17 blood and 2 CSF) with only 1 gastroenteritis (stool) source cases. Isolates from the western region were associated with invasive disease (48 blood and 1 CSF), gastroenteritis (25 stool), and other focal non-invasive infections (5 abscesses/pus and 1 urine) (Table 3). Overall, high serovar diversity was noted. *Salmonella* serovars other than *S*. Enteritidis, *S*. Typhimurium and *S*. Virchow were primarily responsible for gastroenteritis (17/26; 65.4%), whilst *S.* Typhimurium was the leading cause of invasive disease (Table 4). *S.* Typhimurium and *S*. Enteritidis were 5 times and twice as likely to cause invasive disease than gastroenteritis respectively.

**Table 3.**
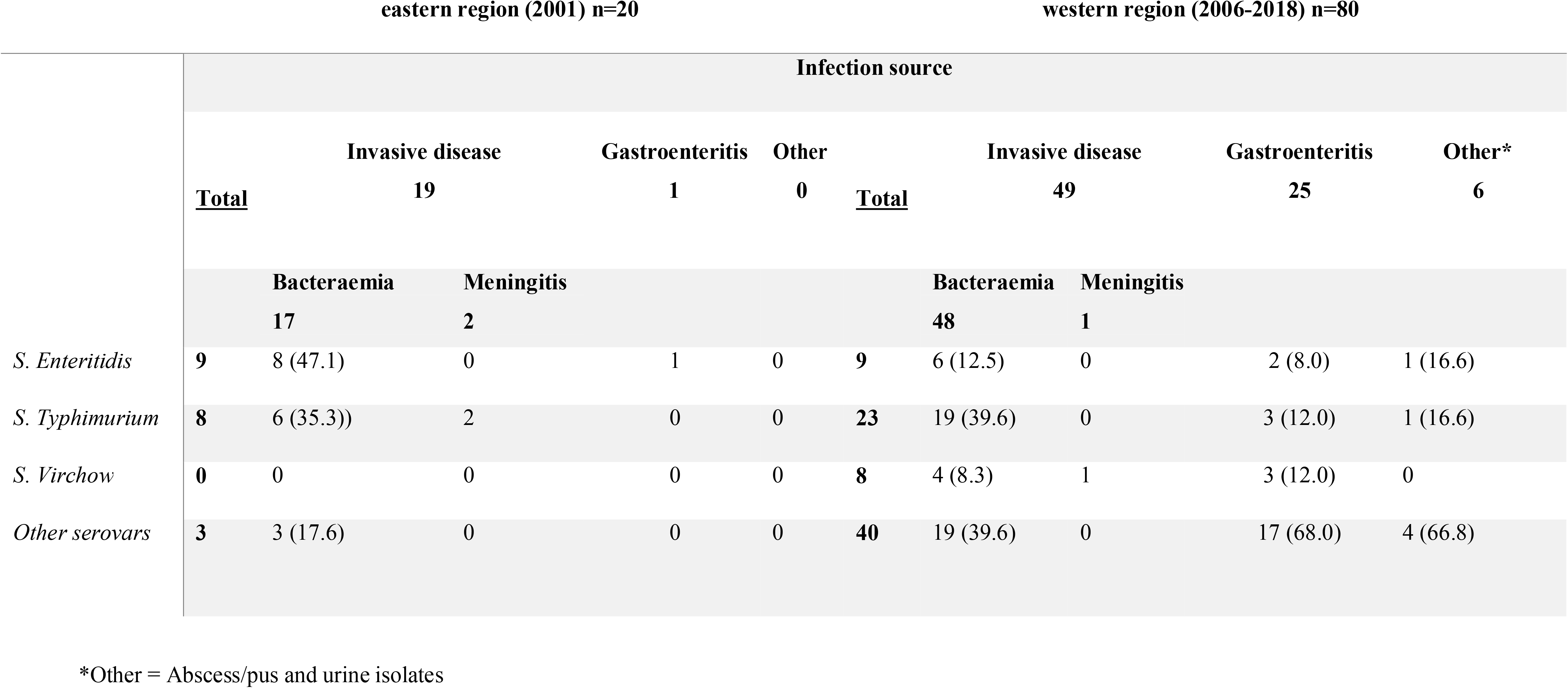
Gambian non-typhoidal *Salmonella* serovar distribution by infection in this study.

**Table 4.** Gambian non-typhoidal *Salmonella* serovar distribution and disease prevalence.

| Serovar | Total | Invasive<br>N (%) | Gastroenteritis<br>N (%) | Others<br>N (%) | Odds of<br>Invasive vs<br>Gastroenteritis | 95% CI | p value |
| --- | --- | --- | --- | --- | --- | --- | --- |
|  | **100** | **68** | **26** | **6** |  |  |  |
| *S. Typhimurium* | 31 | 27 (39.7) | 3 (11.5) | 1 (16.7) | 5.05 | 1.38; 18.48 | **0.014** |
| *S. Enteritidis* | 18 | 14 (20.6) | 3 (11.5) | 1 (16.7) | 1.99 | .52; 7.58 | 0.315 |
| *S. Virchow* | 8 | 5 (7.4) | 3 (11.5) | 0 (0) | 0.61 | 0.13, 2.75 | 0.519 |
| Other serovars | 43 | 22 (32.4) | 17 (65.4) | 4 (66.7) | 0.25 | 0.10; 0.66 | 0.005 |

### Sequence types and eBurst groups

All *S.* Typhimurium were in eBG1 and assigned to a single sequence type, ST19 with one or two allelic variants (Table 5). All *S.* Enteritidis belonged to eBG4 assigned to ST11 and ST1925, including two isolates having single locus allelic variants. *S.* Virchow was in eBG9 and assigned to three different STs as follows: ST181, ST755 and ST841. The four *S.* Hull isolates belonged to eBG330 and assigned ST1996 with single locus variant. *S*. Stanleyville eBG79 (ST339), *S*. Poona eBG46 (ST308) and *S*. Give eBG67 all belonged to ST516.

**Table 5.** eBurst groups of the major Gambian NTS serovars responsible for clinical disease.

| Serovars | Sequence type | eBG | Number |
| --- | --- | --- | --- |
| *S. Enteritidis* | 11 | 4 | 15 |
|  | 1925 | 4 | 1 |
|  | 11 | 4 | 2* |
| *S. Typhimurium* | 19 | 1 | 29 |
|  | 19 | 1 | 2* |
| *S. Virchow* | 181 | 9 | 2 |
|  | 755 | 9 | 1 |
|  | 841 | 9 | 4 |
|  | 841 | 9 | 1* |

\*One or two allelic variants of ST.

### AMR genes, AMR phenotypes, plasmid replicons and virulence genes

Antimicrobial resistance genes belonging to eight classes of antimicrobials were detected, plus a biocide tolerance genetic determinant. Interestingly, genes *aac(6’)-Iaa_1* and *mdf(A)_1* encoding an aminoglycoside modifying enzyme and a multidrug transporter, respectively, were present in all strains, but did not have any detectable phenotypic effects on our isolates (Figure 2). Other AMR genes were harboured by 16/100 (16.0%) isolates and confer resistance to aminoglycosides (*aph_*3_Ib and *aph_*6_Id; n=12), tetracyclines (*tet_*A and *tet_*B; n=4), trimethoprim (*dfr*A14*, dfr*A7 and *dfr*A8; n=8), sulfamethoxazole (*sul2* and *sul1*; n=7), ampicillin (*blaTEM-*1B; n=8), fosfomycin (*fos*A7_1; n=8), azithromycin (*mph_*A; n=3) and chloramphenicol (*cat*A1_1; n=2) (Figure 2). Possession of three or more AMR genes was found in 9/100 (9.0%) isolates, 7/9 (77.8%) of which were found in *S*. Enteritidis.

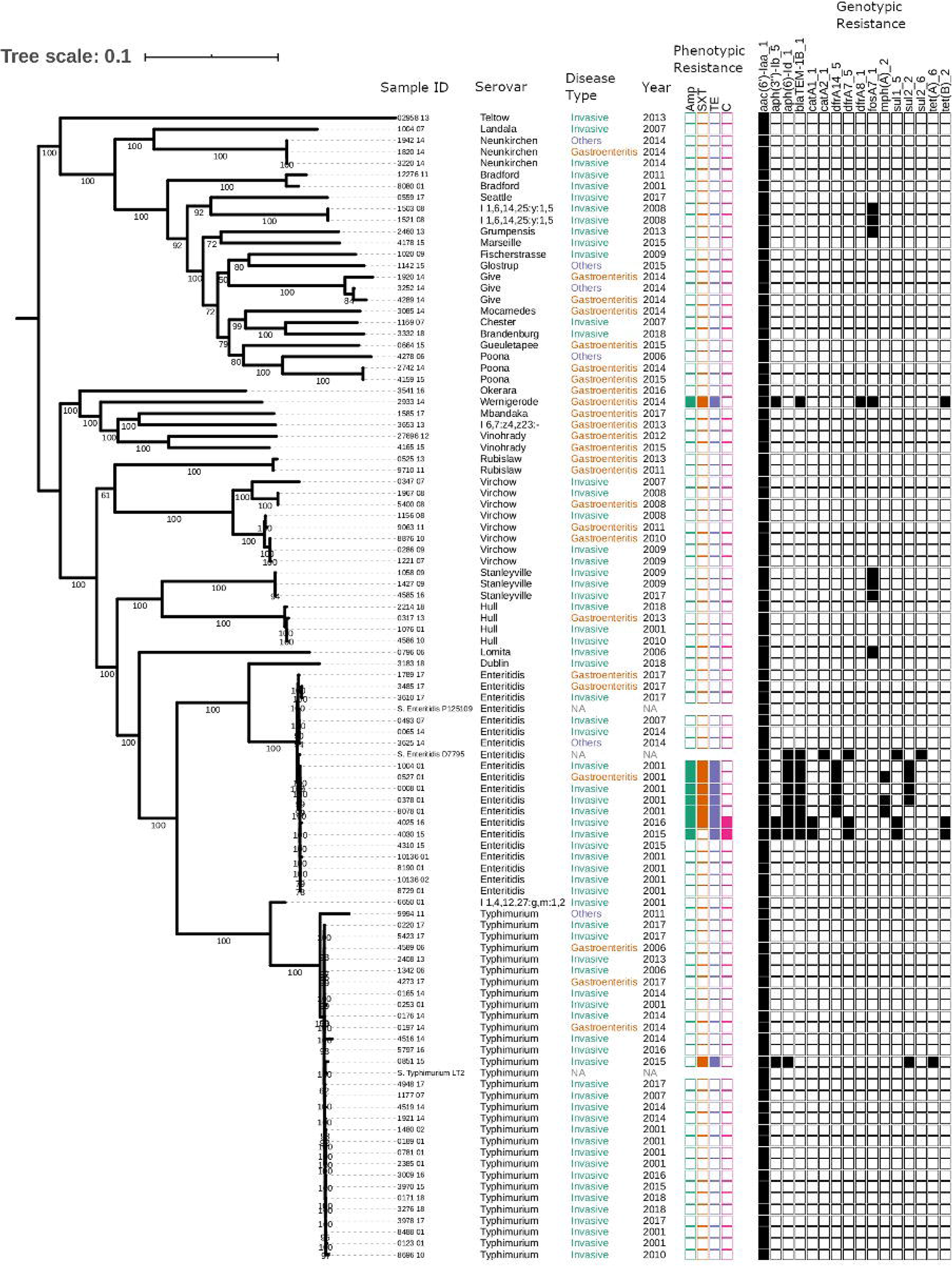

Phenotypic resistance was observed for tetracycline, ampicillin, sulfamethoxazole-trimethoprim and chloramphenicol in 9/100 (9%) isolates, correlating with the presence of resistance genes, except for streptomycin, fosfomycin and azthromycin that were not phenotypically tested (Figure 2). The odds of resistance to ampicillin, sulfamethoxazole-trimethoprim and tetracycline were 51, 20 and 25 times more likely for *S*. Enteritidis than all other serovars combined (Table 6). No resistance to gentamicin was observed phenotypically despite the presence of two aminoglycoside resistance genes (*aph_3_Ib* and *aph_6_Id)*, which only confer resistance to streptomycin.

**Table 6.** Odds of *S.* Enteritidis resistance against other NTS serovars causing disease in The Gambia.

| Antimicrobials | *S. Enteritidis*<br>n=18 | Other serovars<br>n=72 | Odds of *S.*<br>Enteritidis vs all<br>serovars | 95% CI | P-value |
| --- | --- | --- | --- | --- | --- |
| *Ampicillin* | 7 | 1 | 51.5 | 5.78; 459.60 | <0.001 |
| sulfamethoxazole-trimethoprim | 6 | 2 | 20.0 | 3.61; 110.74 | 0.001 |
| Tetracycline | 7 | 2 | 25.5 | 4.68; 138.39 | <0.001 |
| Chloramphenicol | 2 | 0 | 1 | NA | NA |

Nineteen different plasmid replicons were detected in 66/100 (66%) isolates; 7 isolates harboured one plasmid replicon, 39 harboured two, 16 harboured three and 4 harboured four plasmid replicons (Supplementary table 2). The most common plasmid types were *IncFII* (n=55) and *IncFIB* (n=50), harboured by all *S*. Typhimurium and all but the two chloramphenicol-resistant *S.* Enteritidis (Table 7). The *IncN_1* plasmid was associated with MDR including azithromycin resistance and was only found in *S*. Enteritidis from the eastern region (Figure 4a). The *Inc* plasmid is reported to be associated with beta-lactam, streptomycin and sulphonamide resistance. Interestingly, the *IncI*1_Alpha was harboured by the two chloramphenicol MDR *S.* Enteritidis strains from the western region and the susceptible strains from the eastern region (Figure 4b). Notably, no plasmid replicons were detected in *S.* Bradford, *S.* Hull, *S.* Stanleyville, *S.* Rubislaw, *S.* Vinohrady and 1,4,12,27:g,m:1,2 serovars. A total of 115 virulence genes were found with notable absence of *entA, entB, entE, faeC, faeD, faeE, fepC, fepG* in all Gambian NTS serovars (Figure 3). We also identified virulence genes encoding toxins, fimbriae and flagella facilitate invasion, adhesion, type III secretion and survival within host, among other functions increasing bacterial virulence (Figure 3). The highest average number of virulence genes was seen for *S.* Typhimurium (mean: 111/115; 96.52%) with the notable absence of the *cdtB, ssPH1* and *shdA* genes. In contrast, the number of virulence genes *S. Enteritidis* isolates was smaller (mean: 105/115, 91%), with most isolates missing the *cdtB, gogB, grvA, shdA, sinH, slrP, sseK2* and *ssPH1* genes.

**Table 7.**
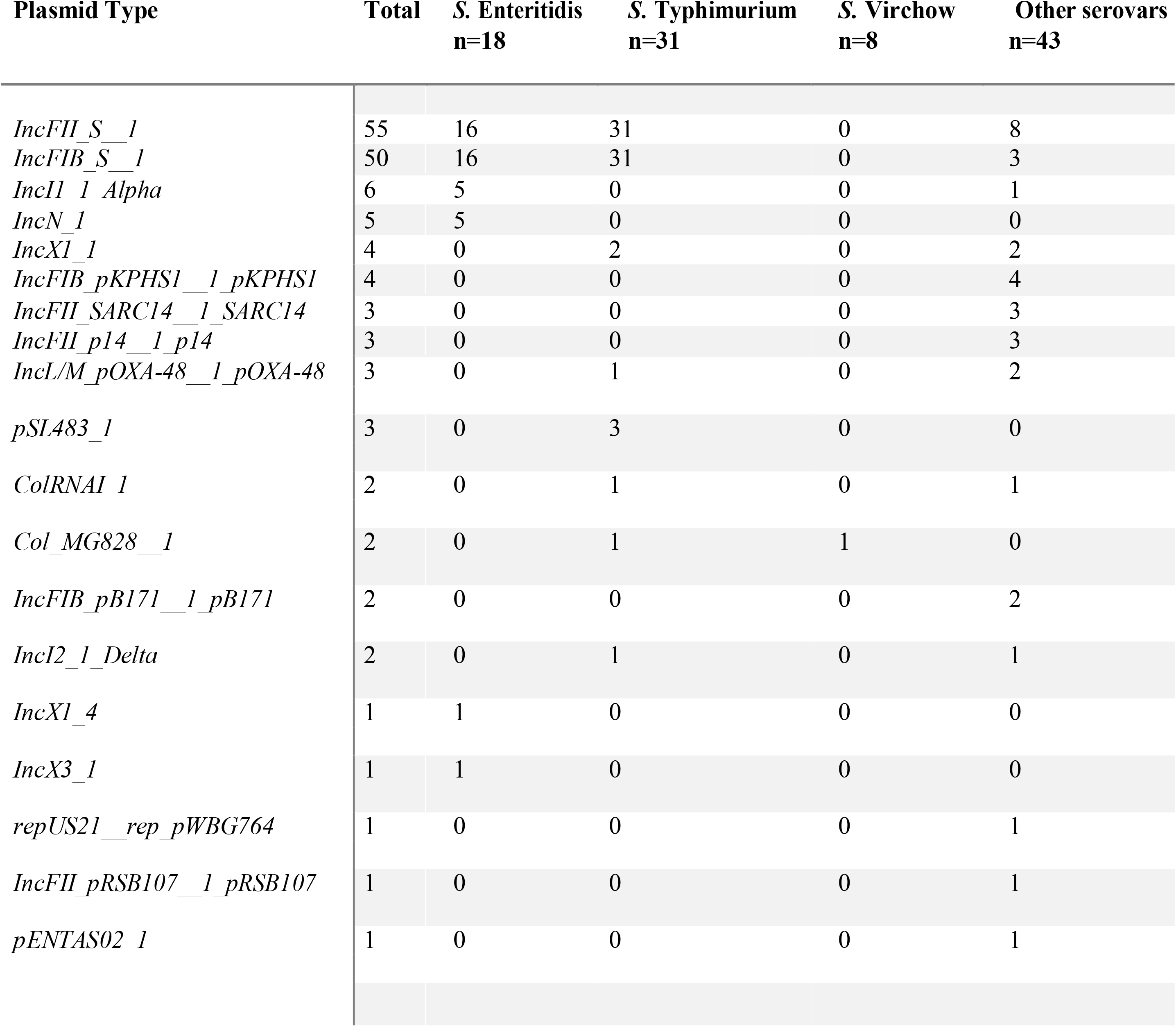
Summary of plasmid replicons and serovar in The Gambian non-typhoidal *Salmonella* isolates.

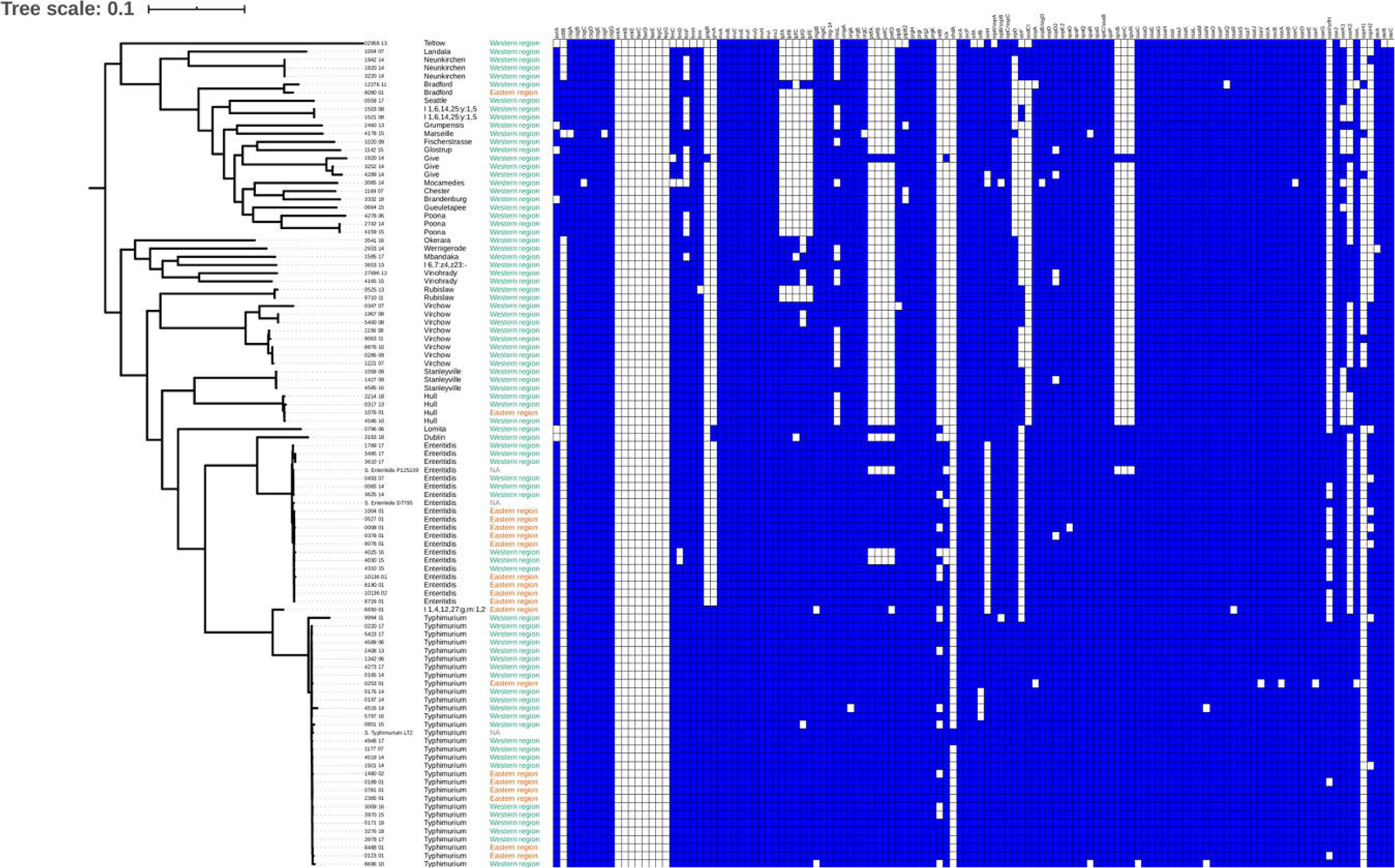

### Phylogenetic analysis

The SNP analysis showed the isolates are clustered into respective serovars and geographic location (Figure 3). *S.* Typhimurium ST19 demonstrasted clonality whilst *S*. Enteritidis was much diverse (Supplemetary figures 1 and 2). To put our data within the wider regional context, we compared our *S*. Enteritidis strains to 495 available African *S*. Enteritidis genomes from the European Nucleotide Archive (ENA). Our analysis revealed considerable genetic diversity which fell into three clades: the North American poultry-associated clade [30] and the global epidemic clade known to cause human gastroenteritidis in addition to the West African clade known to cause invasive diseases carrying the *cat*A1 gene [1] (Figure 4a). All *S*. Enteritidis strains from the eastern region (n=9) and 3 from the western region fell within the West African clade, clustering closely with *S.* Enteritidis strains from Ghana, Guinea and Mali. Among these, 7/12 (58.3%) were MDR (5 from the eastern region and two from the western region) and the remaining were pan susceptible strains (4 from the eastern region plus one western region). Among all strains within the West African clade from the subregion in the study, azithromycin resistance *mph*_A_2 gene was only harboured by strains from the eastern region (Figure 4b). Strains belonging to the North American clade were isolated from blood (n=1) and stool (n=2), whilst those within the global epidemic strains were isolated from blood (n=2) and urine (n=1).

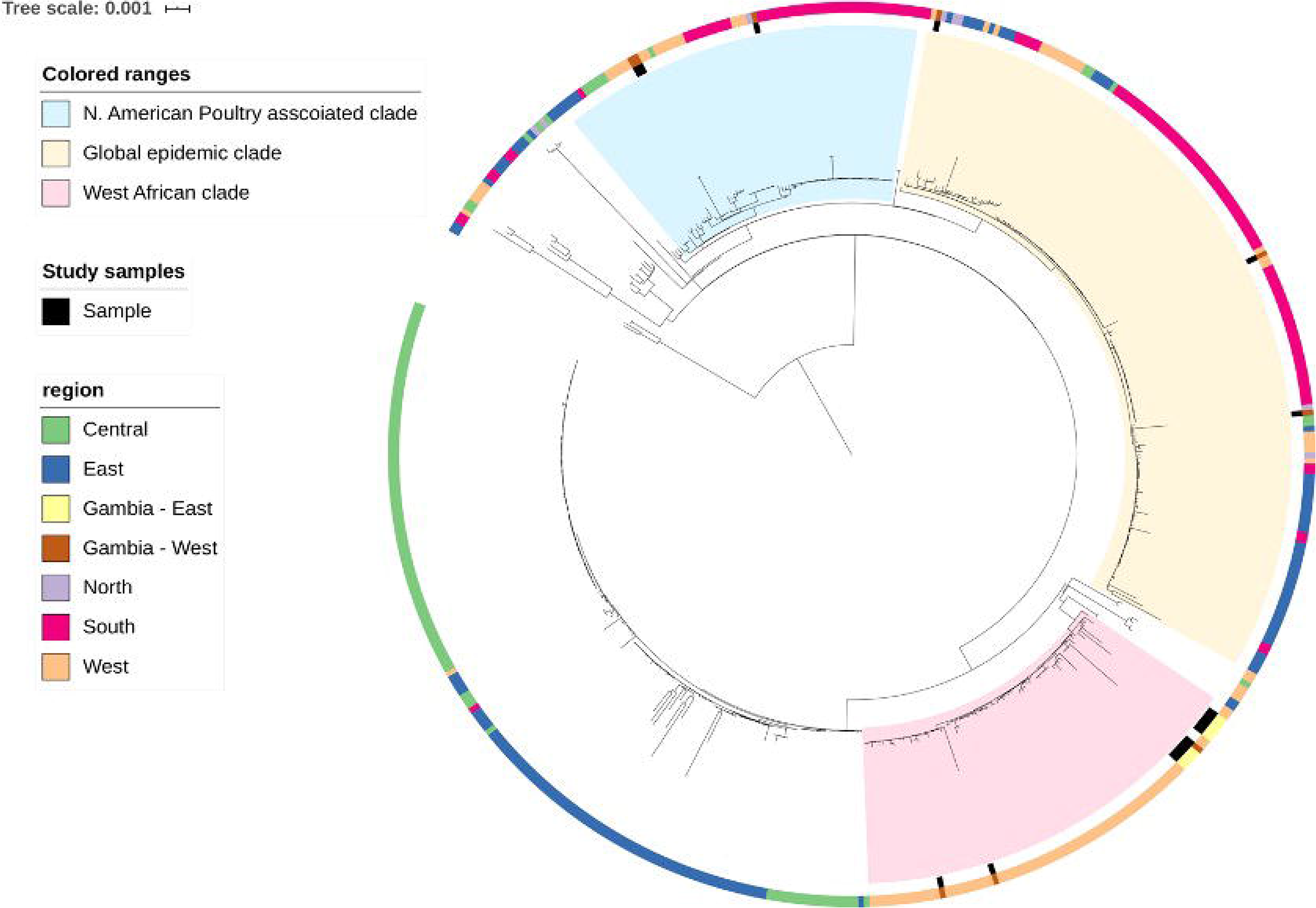

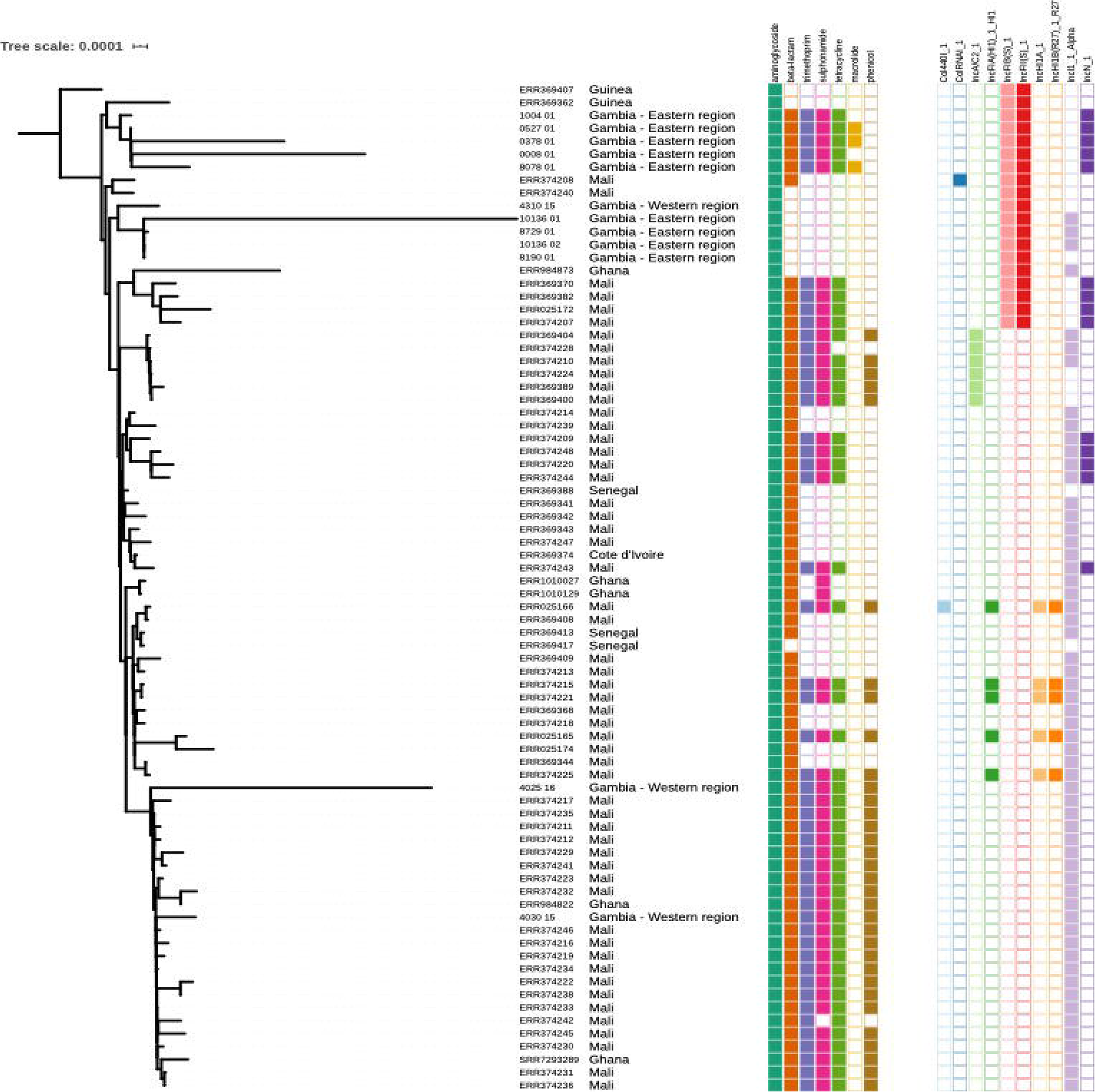

## Discussion

We used phylogenetic analysis to confirm the circulation of a diverse range of NTS serovars including, the epidemic *S*. Enteritidis West African clade mainly in the eastern region of The Gambia, as far back as 2001. This clade is associated with high mortality, harbouring MDR and exhibit genome degradation facilitating an invasive lifestyle, warranting further epidemiological investigations and surveillance as it has important implication in treatment [1,31]. Interestingly, not all *S*. Enteritidis in this study within the clade harboured resistance genes. The *S.* Enteritidis exhibited diversity with other global lineages present. Remarkably, this study found a closely related clonal lineage of *S.* Typhimurium ST19 causing invasive NTS disease as opposed ST313 virulent lineage, which was reported to be circulating in other parts of sSA [7,32]. A possible explanation is that the sequence type ST313 is mainly associated with HIV infection which has a low prevalence in The Gambia [19]. Nevertheless, Panzenhagen *et al.,* reported *S.* Typhimurium ST19-lineage that has evolved in Brazil similar to ST313 restricted to invasive diseases [33]. This unique pathogenesis warrants further comparative genomic and epidemiological investigations into the *S.* Typhimurium ST19-lineage [13].

Two-thirds of serovars responsible for gastroenteritidis in this setting were serovars other than *S*. Typhimurium and *S.* Enteritidis as opposed to other parts of the world where these two serovars account for the highest burden [34]. The great diversity in serovars causing gastroenteritis as opposed to iNTS may suggest that NTS gastroenteritis may not be a predisposition to iNTS in The Gambia. In addition, NTS was not a major cause of gastroenteritis in this setting [35]. Although a huge gap exist regarding transmission dynamics of NTS in sSA [36], more insight is needed in understanding the relationship between NTS gastroenteritis and iNTS disease. In addition, the past two decades has revealed geographical serovar diversity between the two regions, indicating a possible regional specific epidemiological pattern of NTS in The Gambia. However, the time difference in the sampling between the two regions may confound the difference in location and warrant further investigation. Nonetheless, previous phenotypic studies have highlighted these serovar differences [14,15].The changing disease pattern of NTS in sSA, associated with specific lineages of *S*. Typhimurium and *S*. Enteritidis remain a major concern and warrant surveillance [1,9,10,37]. Although a recent decline has been reported for iNTS, it remains a leading cause of bacteraemia in The Gambia [38,39]. Three cases of *Salmonella* bacterial meningitis were included in this study, all of which were found in paediatric patients under 10 years old. Although rare, NTS meningitis has been reported elsewhere in Africa, and is often associated with high case fatality [40,41]. Therefore, NTS needs to be considered in the differential diagnosis of bacterial meningitis following post-vaccine declines in the prevalence of Hib, *Neisseria meningitidis* and pneumococcal meningitis [38,42].

Antimicrobial resistance was found to be correlated with serovar, plasmid replicon and geographical location. MDR was confined within *S.* Enteritidis noted for first-line antibiotics such as ampicillin, sulfamethoxazole-trimethoprim, tetracycline and chloramphenicol. Although no fluroquinolone or cephalosporin resistance was identified, implying these drugs might still be effective in The Gambia, the emergence of azithromycin resistance gene *mph_*A requires further monitoring as a recommended drug of choice for iNTS [43]. Nevertheless, the aminoglycoside resistance gene *aac(6’)-Iaa* and a multi-drug transporter gene *mdf(A)* were present in all serovars including pan-susceptible isolates. This highlights the potential of using genomic-based AMR prediction to monitor AMR determinants for emerging resistance. Our study did not phenotypically test streptomycin susceptibility which lacks clinical breakpoints and it is not used in the treatment of infections. In addition, the streptomycin resistance genes were frequently found to lack expression [44]. While the development of AMR has been mainly attributed to antibiotic misuse in humans and animals, evidence has shown that environmental factors such as poor sanitation, hygiene and access to clean water may be equally responsible for driving resistance in LMIC [45,46].

Although the differences in sampling timepoint was a major limitation in the study, the diversity of AMR between serovars and geographic regions highlights the need for real-time surveillance as well as region-appropriate interventions to effectively combat AMR. Geographic differences seen in AMR may suggest differences in selective pressure and ecological factors thus suggesting need for location specific control measures. The study by Carroll *et al.* underscored distinct factors such as use of antimicrobials in food producing animals as contributing to emergence and dispersal of AMR in humans [47]. Our findings are consistent with other studies that show NTS serovar differences in geographical locations within the same country [47,48]. A correlation between phenotypic and genotypic resistance was also observed (Figure 2). The *IncN* type plasmid was strongly associated with resistance and was found only in MDR *S.* Enteritidis, thus requiring closer surveillance. This plasmid is associated with dissemination of antimicrobial resistance with high potential of spread [49].

We found many virulence genes, however, the identification of virulence factors coding for specific phenotypic traits can be challenging, due to differences in specific traits among serovars [50]. Notwithstanding, the pathogenic success of NTS serovars is directly linked to their plethora of virulence factors aided by host susceptibility, serovar fitness, infectious dose and antimicrobial resistance (AMR) [51]. Further studies are needed to understand the clinical implications of these virulence genes.

There are several limitations in this study. First, the isolates were collected at different time points, with a lag of up to 18 years between the two different regions, which may lead to missing temporal differences. However, the higher AMR prevalence of *S.* Enteritidis in the eastern region as early as 2001 compared to the more recent western region proves the point that AMR emerged a lot earlier and more prevalent in the eastern region. Second, relatively few isolates were analysed from only two regions due to limited microbiology capacity and therefore our results may not reflect the entirety of strains and lineages of the NTS in The Gambia.

In conclusion, this study has revealed great serovar diversity in serovars responsible for gastroenteritis and iNTS and provides evidence for the emergence of MDR *S.* Enteritidis epidemic West African clade in The Gambia. These findings have important implications for antimicrobial prescription policies and regional surveillance of NTS disease. We have demonstrated that a robust genomic epidemiological surveillance of NTS by WGS can be instrumental in generating the critical knowledge and timely information for better disease management and prevention.

## Supporting information

Supplementary figure 1. Phylogenetic tree reconstructed from the pan-genome analysis of S. Typhimurium strains showing plasmids present

Supplementary figure 2. Phylogenetic tree reconstructed from the pan-genome analysis of S. Enteritidis strains showing plasmids present

## Acknowledgement

We thank the following for their support in the study: Jay C.D Hinton, Senior Investigator Award grant recipient of the Liverpool for leading the 10k Salmonella project, Nurudeen Ikumapayi of MRCG@LSHTM for support in providing the isolates from the eastern regions. We are also grateful to Jarrai Manneh and Abdul Karim Sesay for providing support and training with the Illumina sequencing in the MRCG@LSHTM, Buntung Ceesay and Mamadou Jallow of MRCG@LSHTM for contributing to the microbiology laboratory analysis.We thank the patients, Clinical Laboratories, and Services of the Medical Research Council Unit.

## Author notes

Supporting data and protocols are provided as supplementary material and available.

## Author contributions

Conceptualiztion: SD, RSB, MA and BKA.

Laboratory analysis and sequencing of isolates in MRCG: SD

Data transfer of sequence reads into analysis pipeline AW, AK and JP

Processing and sequencing of isolates in the UK, including data transfer. BPS

Data curation: SD, AW, AK

Formal analysis: SD and AKM

Writing original draft: SD

Review and editing: SD, RSB, SY, DN, BPS, BKA, MA

Review of final draft: all authors

Supervision: SK, BKA and MA

Project Administration: SD

