## Supplementary figures and images for "Genomic diversity and antimicrobial resistance among non-typhoidal *Salmonella* associated with human disease in The Gambia"

### Supplementary figure 1. Phylogenetic tree reconstructed from the pan-genome analysis of S. Typhimurium strains showing plasmids present

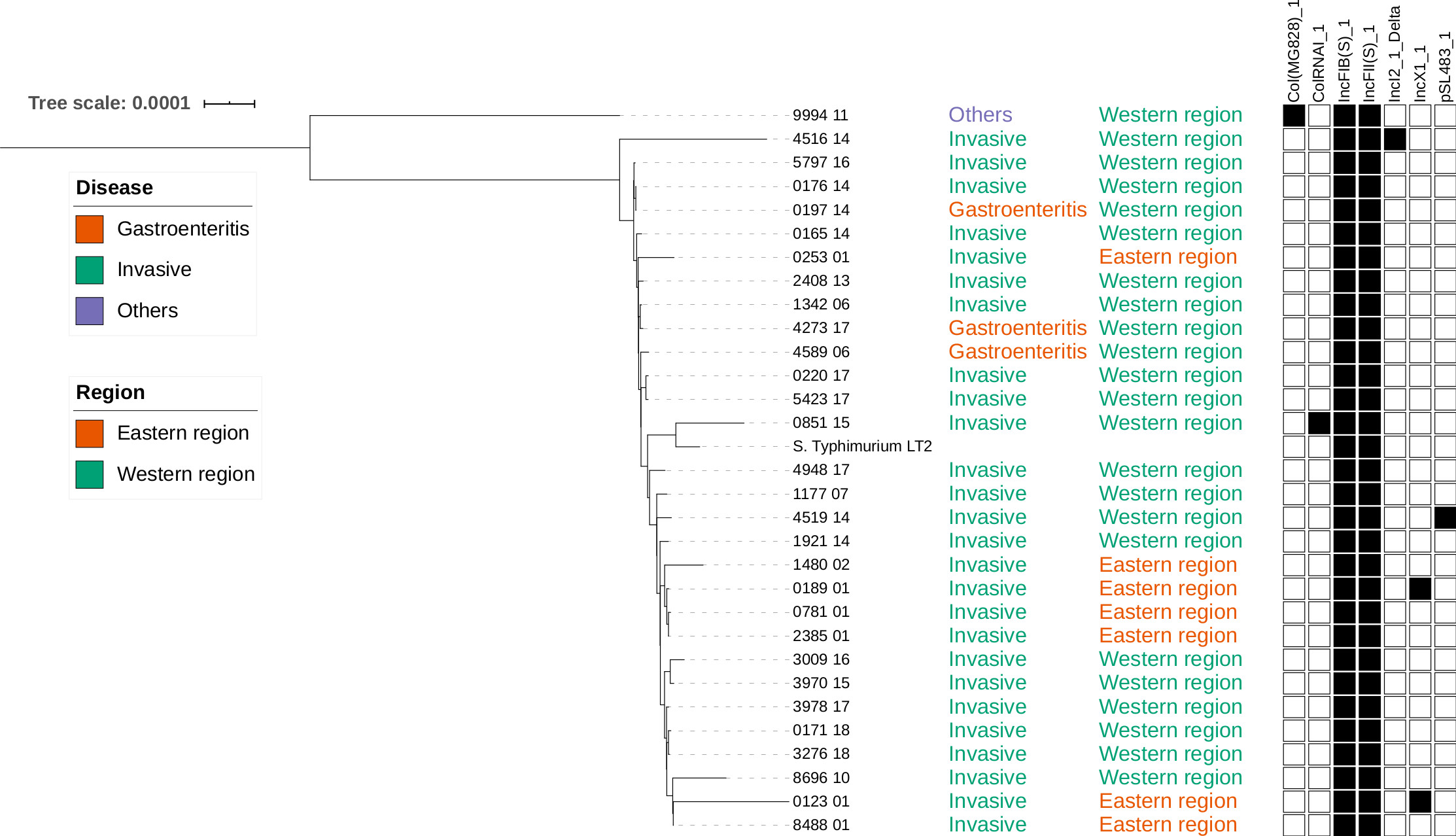

### Supplementary figure 2. Phylogenetic tree reconstructed from the pan-genome analysis of S. Enteritidis strains showing plasmids present

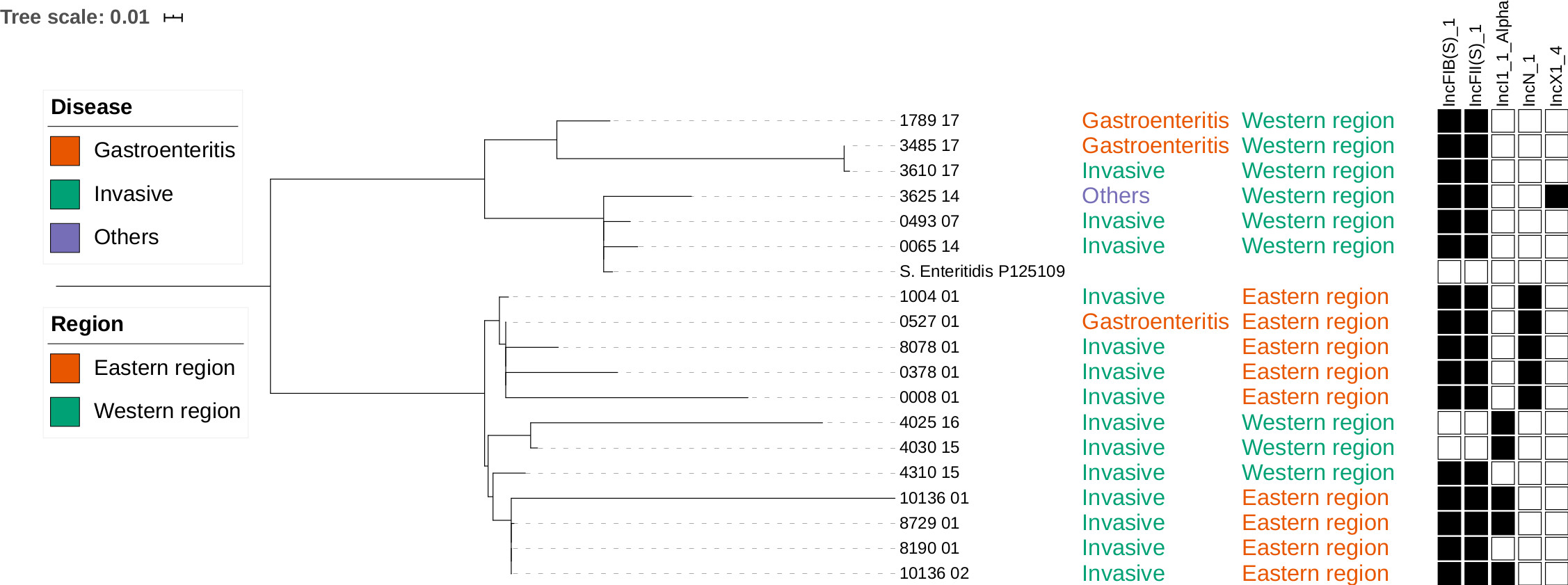
