## Supplementary tables for "Genomic diversity and antimicrobial resistance among non-typhoidal *Salmonella* associated with human disease in The Gambia"

**Supplementary Table 1:** Diversity of other serovars isolated among Gambian NTS in the study, frequency, and associated disease syndrome

| **Other Serovar** | **Number** | **Disease syndrome** |
| --- | --- | --- |
| S. Bradford | 2 | Bacteraemia |
| S. Bradenburg | 1 | Bacteraemia |
| S. Chester | 1 | Bacteraemia |
| S. Dublin | 1 | Bacteraemia |
| S. Fischerstrasse | 1 | Bacteraemia |
| S. Give | 3 | Gastroenteritis, Abscess |
| S. Glostrup | 1 | Abscess |
| S. Grumpensis | 1 | Bacteraemia |
| S. Gueuletapee | 1 | Gastroenteritis |
| S. Hessarek | 1 | Gastroenteritis |
| S. Hull | 4 | Bacteraemia, Gastroenteritis |
| S. Landala | 1 | Bacteraemia |
| S. Lomita | 1 | Bacteraemia |
| S. Marseille | 1 | Bacteraemia |
| S. Mbandaka | 1 | Gastroenteritis |
| S. Mocamedes | 1 | Gastroenteritis |
| S. Neunkirchen | 3 | Abscess, Bacteraemia, Gastroenteritis |
| S. Okerara | 1 | Gastroenteritis |
| S. Poona | 3 | Abscess, Gastroenteritis |
| S. Rubislaw | 2 | Gastroenteritis |
| S. Seatlle | 1 | Bacteraemia |
| S. Stanleyville | 3 | Bacteraemia |
| S. Teltow | 1 | Bacteraemia |
| S. Vinohrady | 2 | Gastroenteritis |
| S. Wernigerode | 1 | Gastroenteritis |
| I 1,4,12,27:g,m:1,2 | 1 | Bacteraemia |
| I 1,6,14,25:y:1,5 | 2 | Bacteraemia |
| I 6,7:z4,z23:- | 1 | Gastroenteritis |

**Supplementary Table2:** Summary of plasmid replicons and serovar harbouring them in Gambian non-typhoidal *Salmonella* isolates

| **Plasmid Type** | **occurrence** | **Serovars harbouring plasmids** |
| --- | --- | --- |
| IncFIB_S__1 | 50 | Enteritidis, Give, Lomita, Typhimurium |
| IncFII_S__1 | 55 | Dublin, Enteritidis, Give, Glostrup, Typhimurium |
| IncFIB_pB171__1_pB171 | 2 | Brandenburg |
| IncN_1 | 5 | Enteritidis |
| IncI1_1_Alpha | 6 | Enteritidis, Poona |
| IncX1_4 | 1 | Enteritidis |
| IncX3_1 | 1 | Enteritidis |
| IncX1_1 | 4 | Dublin, Typhimurium, 1,4,12,27:g,m:1,2 |
| ColRNAI_1 | 2 | Poona, Typhimurium |
| Col_MG828__1 | 2 | Typhimurium, Virchow |
| pSL483_1 | 3 | Typhimurium |
| IncI2_1_Delta | 2 | Poona, Typhimurium |
| IncL/M_pOXA-48__1_pOXA-48 | 3 | Give, Poona, Typhimurium |
| repUS21__rep_pWBG764 | 1 | Chester |
| IncFIB_pKPHS1__1_pKPHS1 | 4 | Give, Neunkirchen |
| IncFII_SARC14__1_SARC14 | 3 | Neunkirchen |
| IncFII_p14__1_p14 | 3 | Neunkirchen |
| IncFII_pRSB107__1_pRSB107 | 1 | Wernigerode |
| pENTAS02_1 | 1 | Teltow |
